# GRU-SCANET: Unleashing the Power of GRU-based Sinusoidal CApture Network for Precision-driven Named Entity Recognition

**DOI:** 10.1101/2024.12.04.626785

**Authors:** Bill Gates Happi Happi, Geraud Fokou Pelap, Danai Symeonidou, Pierre Larmande

## Abstract

**Motivation:** Pre-trained Language Models (PLMs) have achieved remarkable performance across various natural language processing tasks. However, they encounter challenges in biomedical Named Entity Recognition (NER), such as high computational costs and the need for complex fine-tuning. These limitations hinder the efficient recognition of biological entities, especially within specialized corpora. To address these issues, we introduce GRU-SCANET (Gated Recurrent Unit-based Sinusoidal Capture Network), a novel architecture that directly models the relationship between input tokens and entity classes. Our approach offers a computationally efficient alternative for extracting biological entities by capturing contextual dependencies within biomedical texts.

**Results:** GRU-SCANET combines positional encoding, bidirectional GRUs (BiGRUs), an attention-based encoder, and a conditional random field (CRF) decoder to achieve high precision in entity labeling. This design effectively mitigates the challenges posed by unbalanced data across multiple corpora. Our model consistently outperforms leading benchmarks, achieving better performance than BioBERT (8/8 evaluations), PubMedBERT (5/5 evaluations), and the previous state-of-the-art (SOTA) models (8/8 evaluations), including Bern2 (5/5 evaluations). These results highlight the strength of our approach in capturing token-entity relationships more effectively than existing methods, advancing the state of biomedical NER.

## Introduction

Named entity recognition (NER) is pivotal for many natural language processing (NLP) and knowledge acquisition tasks. Typically, the task of NER is to identify real-world entities in an unstructured text using categories such as person, location, organization, time, etc. NER is also needed as an initial step in processes requiring question-answering [1, 2], information retrieval [3, 4], co-reference resolution [5], topic modeling [6, 7]), domain expert assistance [8, 9], to name a few. Machine learning and deep learning play an important role in the biological domains to address various challenges [10]. Recent advances in NLP, such as transformers [11], have significantly improved NER performance, particularly in biomedical contexts, where many entities share similar naming conventions across different species and disciplines [12, 13]. Models like Long Short-Term Memory (LSTM) [14] and Conditional Random Field (CRF) [15] have also greatly improved performance in biomedical NER over the last few years [16, 17, 18].

Additionally, word embeddings such as Word2Vec [19, 20] play a crucial role in capturing token semantics by learning from extensive text corpora such as English Wikipedia, PubMed abstracts, and PMC enabling better contextual understanding and token similarity detection. In addition, these embeddings can automatically detect semantic similarities [21]. For instance, if a model learns the “New York” token in certain contexts and relationships, it can also recognize “NY” as a similarly named entity. However, pre-trained embeddings based on general corpora may reduce the effectiveness of applying NER to domain-specific tasks. This can lead to an information overload for entity recognition within a specific domain, raising concerns about the sensitivity to the quality of embeddings used in this new domain [22]. A recent survey on NER by Maud Ehrmann et al. [23] exposes the problem of lack or imbalanced resources that can influence the capability of the model to perform an accurate prediction.

Therefore, for some architecture, obtaining high-quality embeddings often requires the utilization of large text corpora. For example, BioBERT [24] has been trained on large-scale biomedical corpora comprising approximately 4.5 billion tokens from PubMed abstracts and 13.5 billion tokens from PMC full-text articles. While large-scale language models like BioBERT achieve notable results, their training requires substantial computational resources and fine-tuning, and their performance gains over general NER models can be modest [25]. Furthermore, the reliance on fixed training data introduces biases, as model predictions are influenced by the unbalanced context distributions within the corpora [24].

Although Large Language Models (LLMs) like GPT have revolutionized NLP, their performance in predicting the next token is not without limits [26]. Indeed, their token compression techniques, such as Byte Pair Encoding (BPE), may hide crucial contextual information and introduce bias into the prediction [27], particularly in biomedical NER tasks, where precise token-entity relationships are essential. These biases could result from an unbalanced distribution of contexts in the training data. Additionally, LLMs can struggle with capturing the nuanced relationships between input tokens and output tags, which is necessary for achieving high accuracy in NER [28].

This article presents GRU-SCANET (Gated Recurrent Unit-based Sinusoidal Capture Network), a novel architecture designed to efficiently capture the relationships between input tokens and output tags based on the sentence context. Unlike existing models that rely heavily on large-scale pre-training, GRU-SCANET leverages a lightweight structure utilizing positional encoding to capture token positions, a Bi-directional GRU (BiGru) [29] for learning contextual representations, an attention-based encoder to capture token relations, and a CRF decoder for accurate entity labeling. Our architecture was evaluated on 8 biological NER datasets, demonstrating robust performance across varied corpora and outperforming state-of-the-art models. The results validate the effectiveness of GRU-SCANET in addressing challenges related to unbalanced datasets while maintaining computational efficiency.

The rest of this paper is organized as follows. Section 2 presents the problem formulation. Section 3 presents the GRU-SCANET architecture. Section 4 provides implementation details. Section 5 reports evaluation results and discusses performance compared to other models.

### NER Problem and Formalization

NER models based on BERT or GPT architectures require extremely costly pre-training on corpora and significant processing time for simple tasks to ensure token memorization within various contexts of use [24]. The performance of these models may be degraded if the large corpora on which they rely do not provide a wide enough range or an adequate number of contexts for tokens whose vector representations are critical [30]. For models like BERT and GPT, the absence of context to enrich token representations could make NER processes less effective than expected due to the imbalanced meaningful token representations, appearing in the context to predict the entity type of the next token in the sequence. Thus, we believe it would be preferable to learn how to handle the NER task by directly capturing the relationships between the entities to be detected and their appearances within their contexts to predict them effectively.

Figure 1 represents the four successive steps leading to performing the NER process. (1) Tokenization: The input sentence is meticulously divided into individual tokens, laying the foundation for further analysis. (2) Token Embeddings: Each token embarks on a journey into a high-dimensional vector space, where it is transformed into a unique numerical representation, capturing its semantic essence. (3) Feature Extraction: These token vectors are then skillfully combined to create comprehensive features, meticulously crafted to represent the context of each token within the sentence. (4) Classification: A sophisticated classification model emerges, armed with the extracted features, ready to assign NER tags to each token, identifying the named entities that reside within the sentence.

**Fig. 1.**
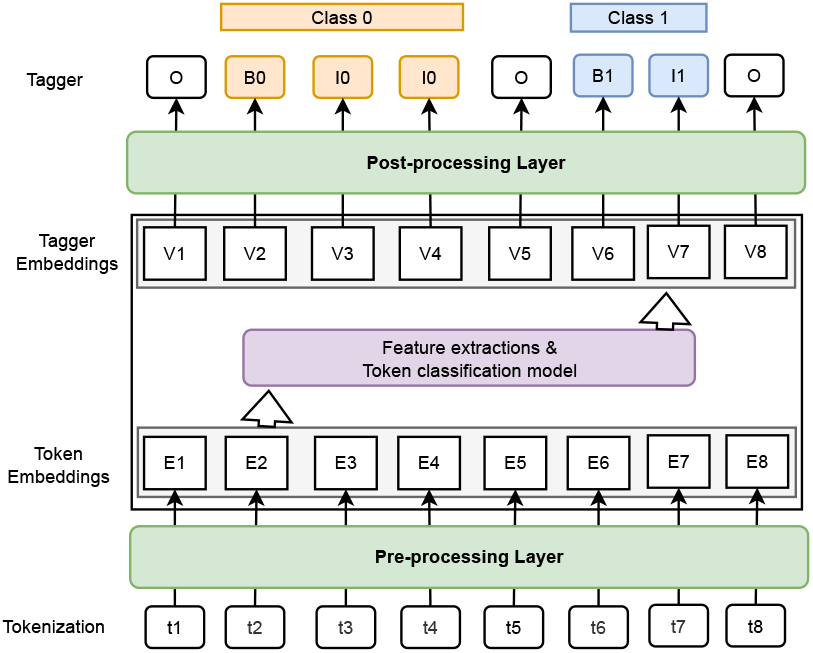
General overview of supervised NER process.

**Fig. 2.**
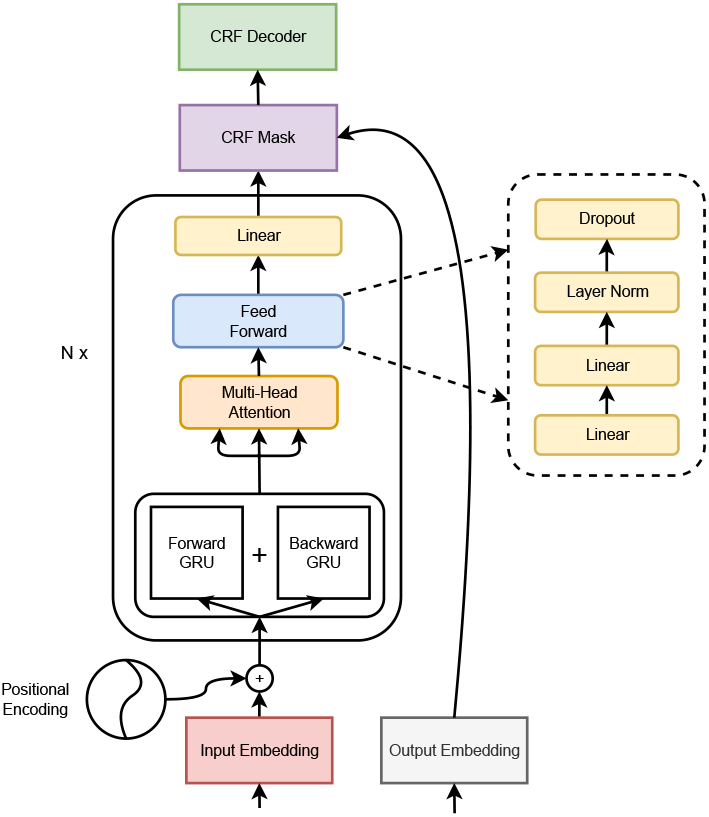
GRU-SCANET architecture.

Consider an input text, which is a sequence of tokens. Each token in this sequence belongs to a token set (*T*) and is associated with a unique index (embedding or order number). Let *X* be a sequence of tokens represented as *X* = [*t*_1_, *t*_2_, *t*_3_, …, *t*_*i*_, …, *t*_*n*_] where *n* is the number of tokens in the sequence (*t*_*i*_ ∈ *T*). Moreover, consider a set of *M* possible entity classes to recognize within *X*. Let us denote as *C*_1_, *C*_2_, …, *C*_*M*_ the *M* possible classes to detect, where *M* is the total number of classes. Each entity class, *C*_*j*_, is associated with three specific tags:

- *B*_*j*_ (Beginning): This tag is used to mark the beginning of a group of tokens that refer to an entity of class *C*_*j*_ in the sequence *X*;
- *I*_*j*_ (Inside): This tag is used to mark a group of tokens in *X* that are part of an entity of class *C*_*j*_, except for the first token;
- *O* (Outside): This tag is used to mark a group of tokens in *X* that are not part of an entity class *C*_*j*_.

The NER problem involves the building of a model (from an architecture) able to associate each token in *X* with one of the three tags (*B*_*j*_, *I*_*j*_, O) of each of the *M* entity classes, to identify and annotate entities within *X*. The *V*_*i*_ of tags (*B*_*j*_, *I*_*j*_, O) are identical on the training data and different during the inference phase due to the probability estimations. In the following sections, we will provide a detailed explanation of each component in our model.

### Approach

GRU-SCANET was trained and evaluated on 8 different biological datasets that were built from the BioCreative benchmarks ^1^. We give the main stages of treatment and their essential points in the following:

1. **Inputs:** from the datasets (test and train from each dataset), we produce a padded *d*-dimensional vector representation of each sentence with their index from the set of words as formalized in section 2 under a form of an integer list. To enhance the high ability of the model to associate word subgroups with their corresponding entity classes, each padded input vector is combined with a positional encoding vector of the same dimension. This addition incorporates the numerical identifier of a word and its position within the input word sequence, effectively specializing the context in which the word is used. This approach significantly improves the distribution of conditional probabilities within the model architecture;
2. **Model Construction:** After forwarding inputs through BiGRU [29] operations, we applied transformation layers. These layers associate the input (word index vector and its positional encoding) with its corresponding output, which is represented as a vector of expected numerical delimiter tag in the output for each entity class (as mentioned in section 2);
3. **Model Training and Testing:** The various datasets are divided into training, development, and test sets. The training step involved optimizing model parameters to minimize prediction errors. There is no need to perform validation steps on development data with GRU-SCANET. Model evaluation was performed on the test set to measure the performance of the model, including precision (P), recall (R), and F1-score (F);
4. **Model Optimization:** by adjusting hyper-parameters such as network size, learning rate, dropout and epochs [29] and weights during processing, we improve the model.

First, the training data were created by matching sequences of words with sequences of tags, using pairs of input words and their corresponding output tags. Next, the model was initialized with an encoder (N=1), which captured the contextual information of the input words. The input sequences were encoded using the *N* encoder (which refers to the transformation layer), and then the CRF (Conditional Random Field) decoding layer was initialized with the expected tags. The training process was repeated over multiple epochs, with adjustment of weights with a back-propagation algorithm, until the model achieved satisfactory performance on the test data. In the following section, we go in-depth into the concepts used by GRU-SCANET.

## Materials and Methods

### Input Embedding and Positional Encoding

Combining sequence embedding with positional encoding aims to consider the relevance of the tokens in their relative position in the sequence *X*, thereby enhancing the model’s ability to capture contextual and spatial information.

- **Input Embedding:** Consider *T* ^*^ being the set of all the sentences. Each sequence of tokens *X* (*X* ∈ *T* ^*^) is represented by matrix denoted *E*(*X*) ∈ ℝ^*d×m*^ (*d, m* ∈ ℕ^*^) in the embedding space. The matrix of *X* is built using the indices of tokens in *T* combined with the function *E*_*i*_. The token embedding function defined as *E*_*i*_ : *T* → ℝ^*m*^. Each token *t*_*i*_ is associated with a *m*-dimensional real-valued vector. *E*(*X*), of the input sequences may not always be the same size as *d*. In such cases, padding is necessary. Padding involves adding a special token with a zero *m*-dimensional vector representation. This ensures that all sequences have a uniform length for further processing. For example, considering that *X* = [*t*_1_, …, *t*_8_] and *d* = 8 according to Figure 1, *E*(*X*) = [*E*_1_(*t*_1_), …, *E*_8_(*t*_8_)] and if *d* = 10, *E*(*X*) = [*E*_1_(*t*_1_), …, *E*_8_(*t*_8_), *E*_*_(0), *E*_*_(0)]. Generally, *d* is defined from the size of the longest sequence of input tokens in the annotated training data and *m* is fixed to an integer constant (e.g.: *m* = 256).
- **Positional encoding:** is typically added to sequence embedding to introduce a notion of position. It uses trigonometric functions to assign specific values to each position in the sequence. A commonly used formula for positional encoding is defined below : Let *t*(∈ ℕ) be the desired position in an input token sentence of size *d. d* represents the max length of input token sequences previously seen. 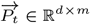 be the corresponding positional encoding of token in position *t* in *X*[11]. *t* belongs to {0, …, *d* −1}, *i*(∈ ℕ) belongs to {0, …, *m*1− 1} and *k* ∈ ℕ.

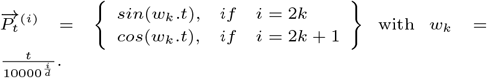

### Encoder stack

#### Forward and Backward GRU models

*Note:* the input of this block is the addition member to member of positional encoding and the embedding of 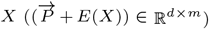.

The Forward and Backward GRU (Gated Recurrent Unit) models are variations of a type of RNN called BiGRU [29]. These models process sequence data, such as sentences or temporal sequences. The Forward GRU model is designed to process sequences chronologically, from left to right. It considers the past information to predict future states. At each time step, the Forward GRU receives the current input and the previous hidden state to generate a new output and update its hidden state. On the other hand, the Backward GRU model processes sequences in reverse order, from right to left. It uses future information to predict past states. At each time step, the Backward GRU receives the current input and the next hidden state to generate new output and update its hidden state. Simultaneously using both the Forward and Backward GRU models enables capturing contextual information from the past and future of a sequence. This allows associating the sequence with its corresponding output class, which is then processed by the subsequent layer to ensure chronological attention.

*Note:* the output of this block is *BiGru*(*E*(*X*)) ∈ ℝ^*d*^.

#### Multi-Head Attention

*Note:* the output of the BiGRU layer is assigned to *Q* = *K* = *V* = *BiGru*(*E*(*X*)).

In the NER context, the multi-head attention mechanism helps to capture relationships and dependencies between tokens in a sequence to represent named entities. The main parameters of the Multi-Head Attention (MHA) process in transformers architectures include the attention heads (default: h=4) having the respective dimensions *d*_*q*_, *d*_*k*_, *d*_*v*_ ∈ ℕ(*d*_*q*_ = *d*_*k*_ = *d*_*v*_ = *d/h* = 256*/*4 = 64, according to 4.1, *d* = 256) for learnable matrices 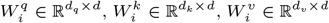.This allows each head to attend to different parts of the input token sequence (from the previous layer) and focus on the non-similar relationships. For each head, the attention scores are calculated by computing the dot product between the query and key representations. The attention scores are then scaled and passed through a softmax function to obtain attention weights. The attention weights compute a weighted sum of the token value representations. This emphasizes the importance of different positions in the input token sequence based on their relevance to the query. The weighted sum is then concatenated across all the heads and projected back to the original dimension using a weight matrix (*W*^*o*^ ∈ ℝ^*h×d*^).

Finally, the output of the MHA process is obtained by applying layer normalization and residual connections to the concatenated and projected representation. This helps stabilize the learning process and allows the model to capture the dependencies and relationships within the input sequence.

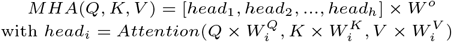

In practical implementation, the attention function is computed on a set of queries packed together into a matrix Q [11]. The keys and values are also packed together into matrices K and V. The output matrix is then computed by applying the attention mechanism:

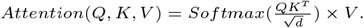

#### Feed Forward Network Layer and Last Linear Transformation

The feedforward sub-block, a crucial component of the Transformer architecture, comprises four sequential linear transformation operations. It processes the output from the BiGRU layer and generates a d-dimensional vector for the final linear layer of the architecture. The first linear layer within this block employs the ReLU [31] activation function. The hidden dimension for all layers within the feedforward sub-block is set to dff=512. The feedforward sub-block consists of four linear transformation operations, each represented by a matrix multiplication and a bias addition.

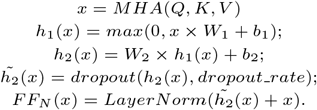

Above *h*_1_, *h*_2_ represents the successive linear layers in the sub-block. *W*_1_ and *W*_2_ ∈ ℝ^*h×d*^. The dropout operation aims to prevent overfitting in neural networks by “dropping out” a certain proportion (dropout rate) of neurons (and their connections) from the network.

Following the feedforward sub-block, a final linearization step is performed using the operation:

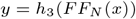

Here, *h*_3_ represents the final linear layer, *FF*_*N*_ (x) denotes the output of the feedforward sub-block, and x represents the input to the sub-block.

### Decoder stack: CRF (Conditional Random Field)

The CRF decoder’s [32] objective is to select the most probable index tag for all the input tokens, considering the input feature scores and index tag transitions. It considers the potential scores of tag sequences generated by the model and the transitions between consecutive index tags. It computes the probability of each index tag sequence using the Viterbi decoding algorithm [33].

- **Output Embedding:** Let *T*_*g*_ = {*B*_*j*_; *I*_*j*_; *O*; 1 ≤ *j* ≤ *M*} be the ordered set of all the taggers of tokens for the *M* classes to detect. Each token *t*_*i*_ in the sequence *X* is associated with a numeric-valued tag *o*_*i*_(*o*_*i*_ ∈ *T*_*g*_) and generates a *d*-dimensional vector *O*_*X*_, which will be utilized by the decoder. *O*_*X*_ has same dimension *d* than *E*(*X*).
- **CRF Mask:** The CRF mask is applied by multiplying the output vector sequence *O*_*X*_ with the binary mask. The masked elements are ignored during the loss calculation and do not contribute to the final predictions. This ensures that only the valid tokens of the sequence are considered during training and prediction, thereby improving the performance and efficiency of the model.

The model is trained in a supervised way by maximizing the probability of correct output tags. This involves jointly optimizing the model weights from the output *y* and the CRF decoder.

The CRF decoder in the architecture allows for the consideration of dependencies between tags, which can improve the coherence of predictions and the overall quality of the model.

## Experiments

In this section, we aim to demonstrate GRU-SCANET’s ability to maintain its effectiveness with consistent performance when scaled up in the NER recognition process. In Section 5.1, we describe the preprocessing of the datasets. Then, in Section 5.2, we describe the execution environment. In Section 5.3, we present our observations on the performance of GRU-SCANET. In Section 5.4, we present an evaluation of the architecture performance scalability in relation to data size and, consequently, model size. Finally, in Section 5.5, we discuss the key differentiations from existing approaches.

### Datasets

We performed a preprocessing step to ensure that the input-output pairs align with the problem’s illustration (see Figure 1), facilitating the model’s learning and prediction during the training and testing phases. The datasets used include NCBI Disease [34], BC5CDR Disease and Drug/Chem [35], BC4CHEMD [36], BC2GM [37], JNLPBA [38], LINNAEUS [39], and Species-800 [40]. Unlike state-of-the-art models like BioBERT [24], which rely on bidirectional encoding, or GPT [26], which uses an auto-regressive process, our architecture does not perform pre-training on diverse corpora to capture token semantics. Instead, it directly maps tokens to the appropriate classes without relying on such extensive pre-training. Our objective is not to predict masked tokens optimally or adapt them to sequences, but rather to capture the relationships between input and output sequences focused on NER datasets. With GRU-SCANET, there is no need to fine-tune the context for each token with new data, preventing the inclusion of irrelevant information that might affect performance. Our model remains flexible when gradually updated with new data, minimizing disruptions compared to models based on token embeddings, which can be sensitive to statistically underrepresented tokens. During the pre-processing stage, we reorganized the datasets into tokenized sentences paired with their corresponding NER tags, simplifying the subsequent training process.

Moreover, we merged all eight data sources into a single dataset to create a Large Language Model (LLM). Subsequently, each of the eight individual datasets was evaluated by using the generated LLMs. This approach ensures that the models are trained and tested globally on the provided high-quality data without focusing on the specified tasks. More details will be described in section 5.4.

### Experimental setup

The experimental setup for the NER process is based on an architecture comprising one encoder and one decoder. The model is trained using a supervised approach, optimizing the parameters with the Adam optimizer to minimize the loss between the predicted and the ground truth tags. The environment contains a GPU partition with 468 GB of RAM and 8 high-performance GPUs (Nvidia v100). Each compute node is equipped with 64 CPU cores, accompanied by dedicated memory of 7.3 GB per core. To achieve optimal convergence, our training process requires only 2 iterations. The entire training process is completed in less than a day. The learning rate is set to 1e-3, and the dropout rate (0.2) is applied to prevent overfitting. The performance of the model is evaluated using appropriate evaluation metrics such as precision(P), recall(R), and F1-score (F). This experimental setup aims to leverage the power of the architecture and the CRF decoder to achieve accurate and robust NER across various biomedical datasets.

Precision (P):

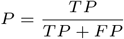

Recall (R):

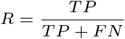

F1-Score (F1):

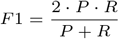

Where:

*TP* - True Positives (correctly predicted positive instances)

*FP* - False Positives (negative instances incorrectly predicted)

*FN* - False Negatives (positive instances incorrectly predicted)

### Overview of results

The GRU-SCANET architecture outperformed the other approaches the evaluated datasets. With a model size of 16 million parameters, GRU-SCANET achieved metrics ranging from 83.52% to 98.64% (Table 3), consistently surpassing state-of-the-art models in various NER tasks. Notably, our architecture outperformed BioBERT in all 8 evaluations and surpassed Bern2 in 5 out of 5 evaluations (Table 3,4,5). Our analysis shows a robust balance between precision, recall, and F1-score, reflecting the model’s ability to accurately label entities in biomedical texts. For example, GRU-SCANET achieved an F1-score of 91.64% on the NCBI Disease dataset and 94.37% on the BC5CDR-chem dataset. These results indicate that GRU-SCANET effectively handles diverse biomedical NER tasks, ensuring high accuracy and minimal error rates.

**Table 1.** Statistics of the biomedical NER datasets.

| Dataset | Entity Type | Number of annotations |
| --- | --- | --- |
| NCBI Disease ( [34]) | Disease | 6881 |
| BC5CDR ( [35]) | Disease | 12 694 |
| BC5CDR ( [35]) | Drug/Chem. | 15 411 |
| BC4CHEMD ( [36]) | Drug/Chem. | 79 842 |
| BC2GM ( [37]) | Drug/Chem. | 20 703 |
| JNLPBA ( [38]) | Gene/Protein | 35 460 |
| LINNAEUS ( [39]) | Gene/Protein | 4077 |
| Species-800 ( [40]) | Species | 3708 |

*Note:* In table 1 The provided information includes the number of annotations from [41] and [42].

**Table 2.** Performance metrics (F1: micro) achieved on benchmark datasets after progressive merging.

| Datasets | Precision | Recall | F1-score | Model’s size |
| --- | --- | --- | --- | --- |
| D1 | 90.21 | 90.21 | 90.21 | 5659818 (6M) |
| D2 | 92.31 | 92.31 | 92.31 | 12133320 (12M) |
| D3 | 92.18 | 92.18 | 92.18 | 12566382 (13M) |
| D4 | 92.88 | 92.88 | 92.88 | 12566940 (13M) |
| D5 | 91.86 | 91.86 | 91.86 | 13039314 (13M) |
| D6 | 92.10 | 92.10 | 92.10 | 14189328 (14M) |
| D7 | 92.68 | 92.68 | 92.68 | 14458198 (14M) |
| D8 | 92.67 | 92.67 | 92.67 | 15079716 (15M) |
| D8 (no MHA) | <u>57.90</u> | <u>57.90</u> | <u>57.90</u> | 10422436 (10M) |

*Note:* Summary table of architecture performance after a progressive increase in data size. We also evaluated GRU-SCANET without the MHA layer that refers to D8 (no MHA).

**Table 3.** Results obtained from the evaluation of the biomedical named entity recognition system.

| Type | Datasets | Metrics | SOTA | BERT | BioBERT v1.0 |  |  | BioBERT v1.1 | GRU-SCANET |
| --- | --- | --- | --- | --- | --- | --- | --- | --- | --- |
|  |  |  |  | (Wiki + Books) | (+ PubMed) | (+ PMC) | (+ PubMed + PMC) | (+ PubMed) |  |
| Disease | NCBI disease | P | 88.30 | 84.12 | 86.76 | 86.16 | <u>89.04</u> | 88.22 | **91.64** |
|  |  | R | 89.00 | 87.19 | 88.02 | 89.48 | <u>89.69</u> | 91.25 | **91.64** |
|  |  | F | 88.60 | 85.63 | 87.38 | 87.79 | <u>89.36</u> | 89.71 | **91.64** |
|  | BC5CDR | P | <u>89.61</u> | 81.97 | 85.80 | 84.67 | 85.86 | 86.47 | **94.25** |
|  |  | R | 83.09 | 82.48 | 86.60 | 85.87 | 87.27 | <u>87.84</u> | **94.25** |
|  |  | F | 86.23 | 82.41 | 86.20 | 85.27 | 86.56 | <u>87.15</u> | **94.25** |
| Drug/Chem | BC5CDR | P | <u>94.26</u> | 90.94 | 92.52 | 92.46 | 93.27 | 93.68 | **94.37** |
|  |  | R | 92.38 | 91.38 | 92.76 | 92.63 | <u>93.61</u> | 93.26 | **94.37** |
|  |  | F | 93.31 | 91.16 | 92.64 | 92.54 | 93.44 | <u>93.47</u> | **94.37** |
|  | BC4CHEMD | P | 92.29 | 91.19 | 91.77 | 91.65 | 92.23 | <u>92.80</u> | **92.83** |
|  |  | R | 90.01 | 88.92 | 90.77 | 90.30 | 90.61 | <u>91.92</u> | **92.83** |
|  |  | F | 91.14 | 90.04 | 91.26 | 90.97 | 91.41 | <u>92.36</u> | **92.83** |
| Gene/Protein | BC2GM | P | 81.81 | 81.17 | 81.72 | 82.86 | <u>85.16</u> | 84.32 | **89.47** |
|  |  | R | 81.57 | 82.42 | 83.38 | 84.21 | 83.65 | <u>85.12</u> | **89.47** |
|  |  | F | 81.69 | 81.79 | 82.54 | 83.53 | 84.40 | <u>84.72</u> | **89.47** |
|  | JNLPBA | P | <u>74.43</u> | 69.57 | 71.11 | 71.17 | 72.68 | 72.24 | **83.52** |
|  |  | R | <u>83.22</u> | 81.20 | 83.11 | 82.76 | 83.21 | **83.56** | 83.52 |
|  |  | F | <u>78.58</u> | 74.94 | 76.65 | 76.53 | <u>77.59</u> | 77.49 | **83.52** |
| Species | LINNAEUS | P | 92.80 | 91.17 | 91.83 | 91.62 | <u>93.84</u> | 90.77 | **98.64** |
|  |  | R | 94.29 | 84.30 | 84.72 | 85.48 | 86.11 | 85.83 | **98.64** |
|  |  | F | <u>93.54</u> | 87.60 | 88.13 | 88.45 | 89.81 | 88.24 | **98.64** |
|  | SPECIES-800 | P | <u>74.34</u> | 69.35 | 70.60 | 71.54 | 72.84 | 72.80 | **95.72** |
|  |  | R | 75.96 | 74.05 | 75.75 | 74.71 | <u>77.97</u> | 75.36 | **95.72** |
|  |  | F | 74.98 | 71.63 | 73.08 | 73.09 | <u>75.31</u> | 74.06 | **95.72** |

*Note:* Following the convention proposed by [24] with their results, we adopt a similar approach to present our summary of results. For each dataset, we provide the Precision (P), Recall (R), and F1-score (F) scores but we use the average micro-metric to evaluate GRU-SCANET. The highest-performing scores are highlighted in bold, while the second-best scores are underlined.

**Table 4.** Evaluations of the biomedical named entity recognition system. (F1 : micro)

| Datasets | BERT |  | RoBERTa | BioBERT | SciBERT |  | ClinicalBERT | BlueBERT | PubMedBERT | GRU-SCANET |
| --- | --- | --- | --- | --- | --- | --- | --- | --- | --- | --- |
|  | uncased | cased | cased | cased | uncased | cased | cased | cased | uncased |  |
| BC5-chem | 89.25 | 89.99 | 89.43 | 92.85 | 92.49 | 92.51 | 90.80 | 91.19 | <u>93.33</u> | **94.37** |
| BC5-disease | 81.44 | 79.92 | 80.65 | 84.70 | 84.54 | 84.70 | 83.04 | 83.69 | <u>85.62</u> | **94.25** |
| NCBI-disease | 85.67 | 85.87 | 86.62 | <u>89.13</u> | 88.10 | 88.25 | 86.32 | 88.04 | 87.82 | **91.64** |
| BC2GM | 80.90 | 81.23 | 80.90 | 83.82 | 83.36 | 83.36 | 81.71 | 81.87 | <u>84.52</u> | **89.47** |
| JNLPBA | 77.69 | 77.51 | 77.86 | 78.55 | 78.68 | 78.51 | 78.07 | 77.71 | <u>79.10</u> | **83.52** |

*Note:* Comparison of GRU-SCANET’s results with those obtained by [43] based on F1-score.

**Table 5.** Performance metrics (F1: micro) achieved on benchmark datasets for biomedical named entity recognition. GSC: GRU-SCANET.

| Datasets | Type | PTC | Hunflair | Bern | Bern2 | GSC |
| --- | --- | --- | --- | --- | --- | --- |
| BC2GM | Gene/protein | 78.8 | 77.9 | 83.4 | 83.7 | **89.47** |
| NCBI-disease | Disease | 81.5 | 85.4 | 88.3 | 88.6 | **91.64** |
| BC4CHEMD | Drug/chemical | 86.7 | 88.9 | 91.2 | 92.8 | **92.83** |
| Linnaeus | Species | 85.6 | 93.2 | 88.0 | 92.7 | **98.64** |
| JNLPBA | DNA | N/A | N/A | N/A | 77.8 | **83.52** |

*Note:* We incorporate our results into the existing version developed by [46] for comparison and further analysis.

A key observation is the equal number of false positives and false negatives across various test instances, leading to similar metrics for precision and recall. This balance highlights the model’s consistency and reliability in tagging entities. For example, if the model is expected to correctly tag 10 entities but makes one incorrect prediction, there will be 1 false positive (FP). Similarly, the number of false negatives (FN) will also be 1, indicating an incorrectly tagged position. When FN equals FP, precision (P) and recall (R) are equal. We calculate the average precision and recall over the entire test dataset, which leads to consistent metrics across different test instances [24]. Furthermore, the ablation study, which involved removing the Multi-Head Attention (MHA) layer, confirmed its crucial role in the model’s performance. The significant drop in F1-score to 57.90% without the MHA layer underscores its importance in capturing complex relationships between tokens. Overall, GRU-SCANET’s stable and high performance across different datasets and experimental setups showcases its capability to address the challenges of biomedical named entity recognition. This architecture sets a new benchmark for NER tasks in the biomedical domain, combining efficiency with accuracy to deliver state-of-the-art results.

### Scalability and Performance

Recall that we conducted evaluations on the following BioCreative datasets (8): BC2GM, BC4CHEMD, BC5CDR-chem, BC5CDR-disease, Corpora, JNLPBA, Linnaeus, NCBI-disease, s800. Using these initial datasets, we created mergers of sets to construct datasets from D1 to D8. For instance, D3 comes from merging the first three datasets listed above. The results are recapitulated in Table 2. In figure 4 and 5, after evaluating each model derived from these merged datasets with the architecture, we observe a stable and slightly increasing model size and stable performances from this architecture. Significant performance fluctuations across the different dataset combinations strongly indicate an imbalance in token distributions between the datasets. Nevertheless, the average value of all metrics is 92.11%. We also performed an ablation study of GRU-SCANET by removing the MHA layer and reevaluating the merged version of the entire dataset (D8). As shown in Table 2 (D8 without MHA), we observe a drastic drop in performance and reduced model size, confirming that the MHA layer is crucial and justifying our performance.

**Fig. 3.**
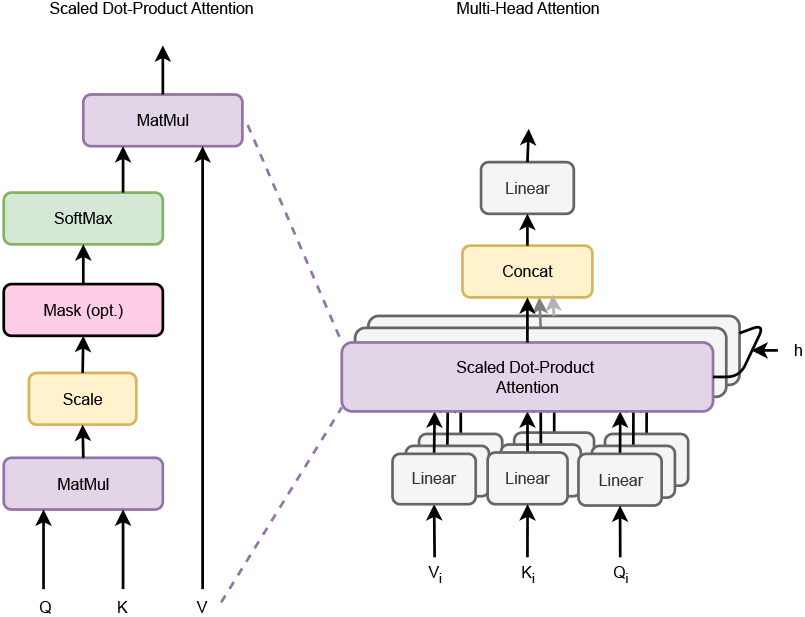
Graphical representation of Multi-Head-Attention [11].

**Fig. 4.**
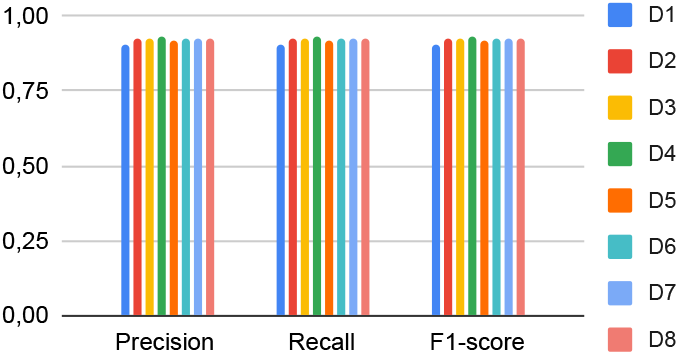
Performance comparison of models of different sizes obtained from the architecture.

**Fig. 5.**
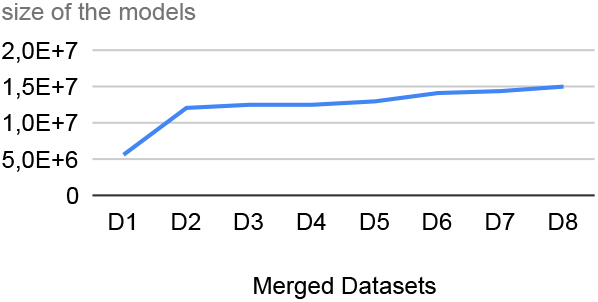
Parameter size curves for different models.

## Discussion

Some studies have shown that it is highly beneficial for auto-regressive models to leverage external corpora tailored to specific domains to optimize token representations to increase the effectiveness in some tasks such as NER [24]. This has been demonstrated by all models, whether they are of biological origin or not, that have adopted the BERT architecture (RoBERTa, BioBERT (cased, v1.0, v1.1), SciBERT, ClinicalBERT, BlueBERT, PubMedBERT, and Bern2). However, in specialized, less-explored domains with a limited training corpus, stabilizing models that aim to optimize token vector representations across multiple contexts can be challenging due to the constraints of the dataset [20]. Our approach GRU-SCANET can maintain high performance without extensive fine-tuning on domain-specific data. This is achieved using BiGRU for contextual learning, multi-head attention for capturing token relationships, and a CRF decoder for precise entity labeling. The ablation study further highlights the crucial role of the multi-head attention mechanism in achieving high accuracy, as evidenced by the significant drop in performance to 57.90% when this component is removed. The scalability of GRU-SCANET is another notable advantage. Our evaluations across progressively larger datasets demonstrate that the model’s performance remains stable and even improves slightly with increased data size. This indicates that GRU-SCANET can effectively handle the growing volume of biomedical literature, making it a robust tool for real-world applications. New approaches have emerged recently to highlight the ability of NER based on LLM prompt contexts with zero-shot or few-shot examples [44, 45], which have so far provided almost no better results despite their varied performance.

## Conclusion

In conclusion, this article introduces an architecture for biological named entity recognition that combines positional encoding, BiGRU, an attention-based encoder, and a CRF decoder. The experimental results validate the effectiveness of this approach in accurately identifying biological entities. Compared to existing models such as Bern, BERT, RoBERTa, BioBERT (cased, v1.0, v1.1), SciBERT, ClinicalBERT, BlueBERT, PubMedBERT, and Bern2, GRU-SCANET offers enhanced accuracy and efficiency in extracting named entities from biomedical texts. Evaluation results on benchmark datasets demonstrate the effectiveness of GRU-SCANET in recognizing various types of entities, including genes/proteins, diseases, drugs/chemicals, and species. Local installation options make GRU-SCANET easily accessible for integration into other systems. GRU-SCANET provides researchers and practitioners a reliable tool for improving biomedical text-mining tasks. Overall, GRU-SCANET holds great potential in advancing research and applications in biomedicine. Future perspectives for this approach include exploring larger and more diverse biomedical datasets, as well as incorporating domain-specific knowledge and ontologies to improve performance in capturing specific biomedical entity types and relationships.

## Availability and implementation

source code at https://github.com/ANR-DIG-AI/GRU-SCANET

## Author contributions statement

B.G. conceived the architecture and conducted the experiments, P.L. conducted the literature reviews, and D.S. and G.F. analyzed the results. We all reviewed the manuscript.

## Acknowledgments

This work was granted access to the HPC resources of IDRIS under the allocation 2024-AD011012511R3 made by GENCI. The authors thank the IRD supercomputing infrastructure, I-Trop, and its staff for their support. The authors thank the South Green Bioinformatics Platform for its IT support.

- **Bill Gates Happi Happi** is a dedicated Ph.D. student in computer science at the University of Montpellier. He works at the IRD and is actively involved in scientific exploration mainly in Entity recognition for knowledge graph augmentation.
- **Géraud Fokou Pelap** is Semantic Web Expert and AI Consultant for M2IA Company. He is also associate researcher in computer science at the University of Dschang in URIFIA Laboratory.
- **Danai Symeonidou** is a researcher in computer science at the French Research Institute for Agriculture, Food and the Environment (INRAE).
- **Pierre Larmande** is a senior researcher in computer science and bioinformatics at the French Institute of Research for Sustainable Development (IRD).

## Footnotes

1 https://biocreative.bioinformatics.udel.edu/

